# Instabilities and Spatiotemporal Dynamics of Active Elastic Filaments

**DOI:** 10.1101/725283

**Authors:** Yaouen Fily, Priya Subramanian, Tobias M. Schneider, Raghunath Chelakkot, Arvind Gopinath

## Abstract

Biological filaments driven by molecular motors tend to experience tangential propulsive forces also known as active follower forces. When such a filament encounters an obstacle, it deforms, which reorients its follower forces and alters its entire motion. If the filament pushes a cargo, the friction on the cargo can be enough to deform the filament, thus affecting the transport properties of the cargo. Motivated by cytoskeletal filament motility assays, we study the dynamic buckling instabilities of a two-dimensional slender elastic filament driven through a dissipative medium by tangential propulsive forces in the presence of obstacles or cargo. We observe two distinct instabilities. When the filament’s head is pinned or experiences significant translational but little rotational drag from its cargo, it buckles into a steadily rotating coiled state. When it is clamped or experiences both significant translational and rotational drag from its cargo, it buckles into a periodically beating, overall translating state. Using minimal analytically tractable models, linear stability theory, and fully non-linear computations, we study the onset of each buckling instability, characterize each buckled state, and map out the phase diagram of the system. Finally, we use particle-based Brownian dynamics simulations to show our main results are robust to moderate noise and steric repulsion. Overall, our results provide a unified framework to understand the dynamics of tangentially propelled filaments and filament-cargo assemblies.

## 1 Introduction

Actin filaments and microtubules are some of the fundamental building blocks of biological systems. The mechanical properties of biological objects often come down to the way those elastic filaments move and bend under external loads or internal molecular motor forces ^1^. Examples include the buckling of cytoskeletal filaments by molecular motors ^2–9^, the sliding deformations of microtubule doublets induced by ATPase dynein in eukaryotic cilia ^10–12^, and the looping of DNA that is crucial to the normal functioning of the cell. In the synthetic realm, filaments made of connected magnetically responsive colloids driven by external fields ^13–16^ or of tailored connected Janus particles ^14^ may also bend or buckle as they move.

From a theoretical perspective, the elastic properties of a thin composite filament can be captured by an elastic line model whose only free parameter is the filament’s bending rigidity. In contrast, modeling the deforming forces requires knowledge of their direction and spatiotemporal variations. In this paper, we focus on the case of follower forces, i.e., forces that always act along the filament’s tangent. One possible realization of this problem is a chain of self-propelled Janus colloids bonding to each other along the same direction in which they self-propel. The best studied example of follower forces at the micro-scale, however, is gliding motility assays, wherein cytoskeletal filaments (actin filaments or microtubules) glide over a surface under the propelling action of molecular motors (dynein, kinesin, or myosin) grafted onto the surface ^17–23^. The interaction between motors and filaments naturally generates tangential forces distributed along the filament with a consistent direction (always towards the same end of the filament). In the absence of obstacles or other restrictions to its motion, such a filament moves in a straight line as if on a molecular-motor powered conveyor belt. This motion, however, is easily disrupted, e.g., by optical trapping, or by surface defects. The motor forces then become compressive and can buckle the filament (see figure 1). Experiments reveal the resulting buckled shape is determined by the strength and type of surface defect or pinning ^17,22^. Buckling instabilities have also been observed in simulations of a variety of models of pinned or clamped motility assays filaments ^22,24–27^. Recent Brownian dynamics simulations also found buckling instabilities in follower-force propelled filaments pushing a cargo ^28^.

**Fig. 1.**
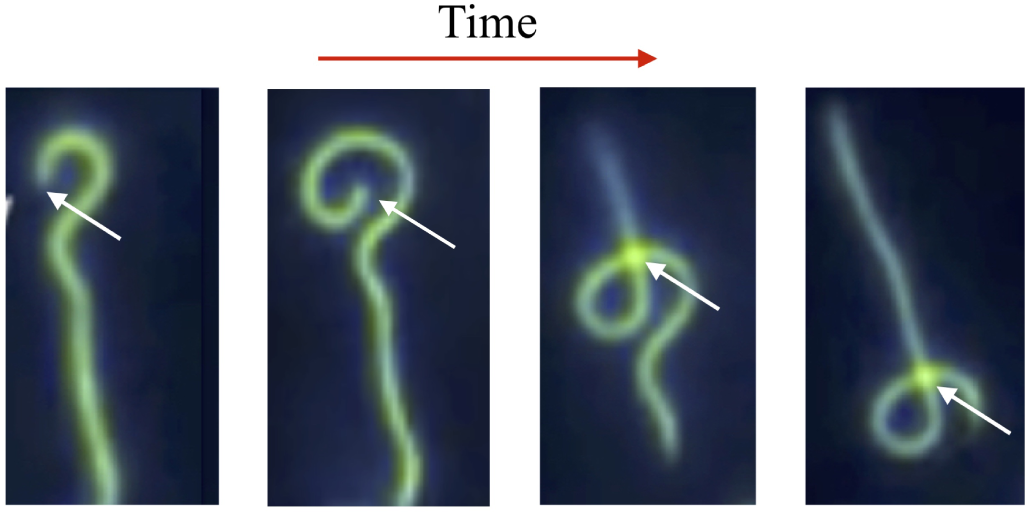
Fluorescence enhanced two dimensional gliding motility assay images showing a microtubule, propelled by kinesin-1 motors, buckling upon encountering a surface defect or other pinning site ^17,41^. The pinning site’s location, indicated by the white arrow in each tile, is fixed, however the field of view changes to follow the filament. (Tile 1) The front of the filament has hit the defect and started to buckle into a near-circular shape. (Tile 2) The pinned, buckled, front end of the filament rotates around the defect. (Tile 3) The front of the filament escapes the defect and resumes straight motion. A region of high curvature persists near the defect as the filament slides through it, suggesting the defect still acts on the filament. At the defect the filament breaks the two-dimensional confinement to slide over itself. (Tile 4) The filament continues its motion through and away from the defect.

Outside of biology, the follower-force induced buckling problem has received significant attention from the engineering community, albeit in situations where inertia plays a role, e.g., water bearing flexible pipes or propelled structures like rockets ^29–39^. Theoretical studies from this body of literature emphasize that tangential forces, being non-conservative, do not lend themselves to the usual energy minimization approaches to buckling ^40^. In order to analyze the stability of the filament, one must study the full dynamical response.

Here, we show that the overdamped version of the follower-force induced buckling problem, exemplified by gliding motility assays, can be similarly analyzed by studying the time-dependent response of the filament to deformations. Combining concepts from classical Euler beam theory, viscous resistive force theory and a coarse-grained description of the motor generated follower forces, we propose a two-dimensional continuum model that governs the shape of actively deformed filaments. This in turn provides us with a unified framework to understand the planar buckling instabilities of biological filaments under follower load and the role of boundary conditions in selecting which instability develops.

The paper is organized as follows. In section 2 we discuss the model and the approximations that lead to it in the context of motility assays. In section 3 we look at the buckling of filaments whose head is held fixed, either pinned (free to rotate) or clamped (unable to rotate). We identify equilibrium base states, the critical points at which they become unstable, and track the non-linear non-equilibrium states that bifurcate therein. We observe that initially straight pinned filaments buckle into a steadily rotating coiled state whereas clamped filaments buckle into a periodic beating state wherein deformation waves travel through the filament. We further identify a simpler model with the same stability phenomenology but whose linear stability is analytically tractable, thus providing additional insight into the full model’s paths to instability. Section 4 extends our linear and non-linear analysis to filament-cargo assemblies. We find that filament-cargo behaviors can be largely understood in terms of the pinned and clamped cases above, with the cargo playing a role similar to either a partial pinning or a partial clamping site depending on its translational and rotational drag coefficients. In section 5 we show that the results of our stability analysis and noiseless non-linear simulations compare well with Brownian dynamics simulations of noisy animated filaments. Finally in section 6 we discuss our results’ place amongst the recent literature on biologically motivated follower-force buckling problems.

## 2 Continuum model for filament dynamics

### 2.1 Geometry, forces and torques

Gliding assays typically feature polar filaments moving in a thin fluid film over a flat surface. Thus we restrict ourselves to planar configurations and model overdamped filaments moving in the *xy* plane with cartesian basis (**e**_*x*_, **e**_*y*_) (see Figure 2). The filament is treated as a slender column with circular section, length *ℓ*, and diameter *σ* ≪ *ℓ*. The material comprising the filament is assumed be a linear elastic material with Young’s modulus *E* and bending modulus *κ* ∼ *Eσ*^4^. For high aspect ratio filaments *ℓ/σ* ≫ 1 as is appropriate in our case, the filament accommodates compressive forces by bending rather than stretching. We therefore treat the filament as inextensible. One end of the filament, called its *head*, is either pinned, clamped, or rigidly attached to a viscous load. The other end, called the *tail*, is free.

**Fig. 2.**
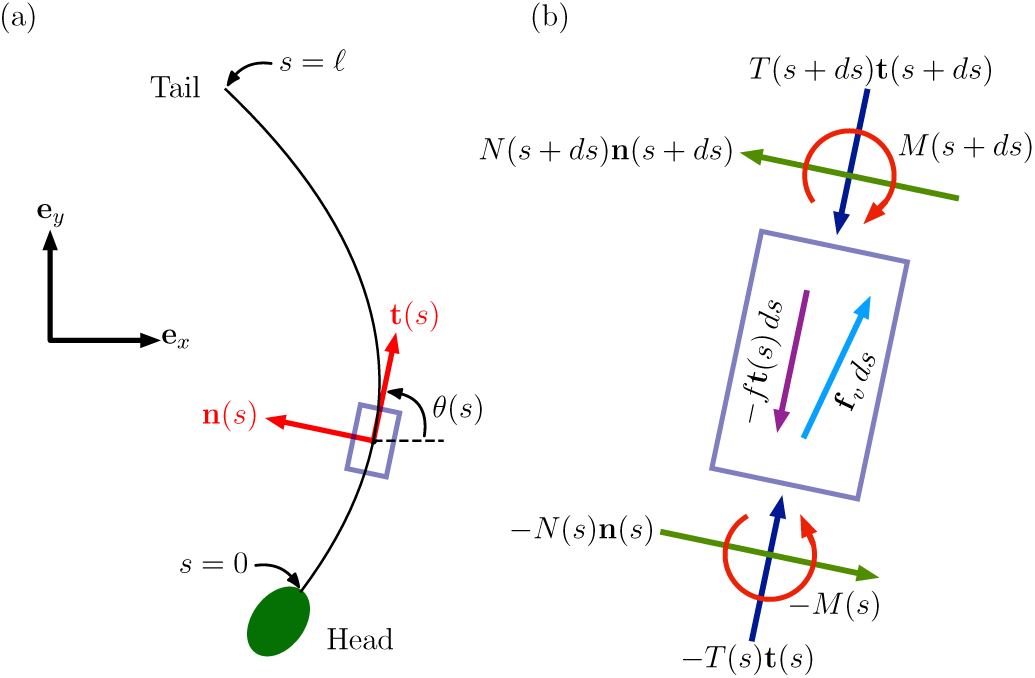
Definition sketches used in the model for the deforming filament. (a) The active force density − *f* **t**(*s*) acts anti-parallel to the local tangent vector **t**(*s*) with *s*, the arc-length, measured from the head. The slender filament is free at *s* = *ℓ* while the end *s* = 0 is constrained in some manner. The filament moves in the two dimensional plane spanned by the tangent and the normal coinciding with the *x* − *y* plane. (b) Sketch of the forces and torques acting on a small differential element *ds* of the filament about location *s*. Shown are the normal *N*(*s, t*) and tangential *T* (*s, t*) components of the cross-sectionally averaged force resultant, the moment (the internal torque), *M*(*s, t*), and the externally acting forces with active (*f*) and dissipative (*f*_v_) components.

The shape and location of the thin filament are described by the parametrization **r**(*s, t*) of its center-line where 0 ≤ *s* ≤ *ℓ* is the arclength along the filament. Since the filament only moves in two dimensions, its shape is fully determined by the angle *θ* (*s, t*) between the positive *x* axis and the filament’s tangent vector at *s*. The unit tangent and normal vectors at point **r**(*s, t*) are then given by **t** = cos *θ* **e**_*x*_ + sin *θ* **e**_*y*_ and **n** = − sin *θ* **e**_*x*_ + cos *θ* **e**_*y*_, respectively.

Force and torque balances demand that internal force and torque densities due to the bending of the filament balance external force densities as illustrated in Figure 2(b). In addition to internal forces and torques, each material point along the filament experiences two external forces: the active motor force, and the dissipative viscous force, both of which can be treated as externally applied force densities.

The active forces are created by regularly spaced molecular motors grafted onto the surface beneath the filament with areal density *ρ*_*m*_. The filament only interacts with motors located within a short distance *δ*_*m*_ of its center-line. Those motors spend a fraction *r* of their time attached to the filament, during which they exert a tangential force with magnitude *F*_*m*_. Unattached or out-of-range motors exert no force. At high motor densities *ρ*_*m*_ and duty ratios *r*, the mean effective force per motor is nearly independent of the motor density ^5^. Assuming further that attached motors are synchronized, and ignoring variations in the magnitude and direction of the motor generated forces ^42^, the effective net force density acting on the filament is − *f* **t** with constant magnitude *f* ∼ *F*_*m*_*ρ*_*m*_*δ*_*m*_*r*.

Passive dissipation comes from the filament’s interaction with the surrounding fluid film. It is a very low Reynolds number environment, meaning inertial effects are negligible. The solid surface beneath the fluid film acts as a momentum sink that screens hydrodynamic interactions, thus we also neglect nonlocal hydrodynamic effects, only keeping local hydrodynamic drag. Let 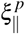 and 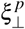 be the corresponding viscous resistances per unit Length in the tangential and normal directions, respectively. In a 3D Newtonian fluid of viscosity *µ*, local resistivity theory predicts 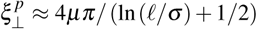 and 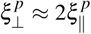. The relationship 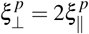 remains true in two dimensional settings with filament deformations confined to a plane^13^. In structured media such as gels, on the other hand, the ratio between the two coefficients may be different ^2,8^.

In addition to this passive hydrodynamic force, dissipative effects also arise from active motor kinetics especially at high duty ratio due to the energy dissipated as attached motors detach. Internal motor friction may also be induced as attached motors are swept by the filament resulting in dissipative forces acting at every location on the filament. We incorporate these motor-induced resistances using the tangential and normal active resistances per unit length 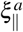 and 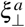. Combining the active and passive contributions, the total drag force density is 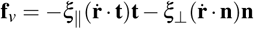 where 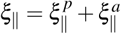 and 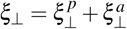are the effective tangential and normal resistances per unit length.

A range of possible viscous effects may be investigated conveniently by treating the *viscosity contrast γ* ≡ *ξ*_⊥_*/ξ*_∥_ as an independent parameter. When motor friction is small such that friction is entirely due to the surrounding Newtonian fluid, then *γ* ≈ 2. If the surrounding fluid instead resembles a freely draining polymer solution (e.g., for very dilute assays), *γ* ≈ 1. For a gel-like material, *γ* ≫ 1. When motor friction is significant, such as for densely packed motors with high duty-ratio, *ξ* _∥_ ≫ *ξ* _⊥_ thus *γ* ≪ 1.

The equations governing the spatiotemporal dynamics of the filament illustrated in Figures 2(a-b) can be derived by invoking force and torque balances (ESM §I). We work in a dimensionless setting by scaling all forces by *κ/ℓ*^2^, lengths by *ℓ*, and times by *ℓ*^4^*γξ* _∥_*/κ*. The cross-sectionally averaged internal force resultant **F** evaluated at **r**(*s, t*) can be decomposed into its tangential *T* and normal *N* components, **F** = *T* **t** + *N***n**. Using this decomposition, we first combine force and torque balances at a representative material point with constraint equations that relate the shape of the filament to the velocity of its centerline, then use the Frenet-Serret relationships to eliminate the state variable *N* in favor of the shape variable *θ* to obtain the two dimensionless coupled partial differential equations

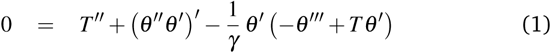

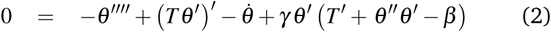

where we have denoted *∂* **r***/∂t* by 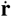 and derivatives with respect to arc-length *s* by primes. Equations (1) and (2) feature two dimensionless parameters:

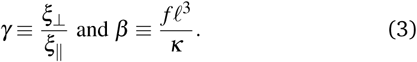

We have already discussed the viscosity contrast *γ*. The activity parameter *β* captures the relative importance of the active force, which causes buckling, to the filament stiffness, which curbs it. One may also write *β* = (*ℓ/*λ**)^3^ where 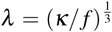 is the length over which compression is accommodated. Thus *β* ≫ 1 corresponds to a very active, very soft, or very long (*ℓ ≫ λ*) filament, whereas *β* ≪ 1 corresponds to a very weakly active, very stiff, or very short (*ℓ* ≪ *λ*) filament.

The range of *β* values accessible to motility assay experiments is remarkably large. The lower bound is zero, obtained by decreasing the motor density or the ATP concentration to zero. The upper bound can be estimated from the values of *F*_*m*_, *ρ*_*m*_, *δ*_*m*_, *r, κ*, and *ℓ* found in the experimental literature. For microtubules driven by kinesin, Bourdieu *et al.* ^22^ *report δ*_*m*_ ∼ 20 nm, *ρ*_*m*_ ∼ 2000 − 5000 *µ*m^−2^, *r* ∼ 1, *F*_*m*_ ∼ 2 − 5 pN, and a persistence length *ℓ*_*p*_ ∼ 5mm at room temperature. Thus *f* may be as large as ∼ 5 *×* 10^−4^ Nm^−1^, and *κ* ∼ 2 *×* 10^−23^ N.m^2^ (consistent with ^2^). Average microtubule lengths of 5 − 10 *µ*m seem common ^1,2,17^, which yields the upper bound *β*_max_ ∼ 3 *×* 10^4^. On the other hand, microtubules can have a broad distribution of lengths with small amounts of longer filaments, up to at least30 *µ*m ^17^, mixed with shorter ones. Since *β* is proportional to *ℓ*^3^, this yields a significantly higher upper bound *β*_max_ ∼ 7 *×* 10^5^. For actin driven by myosin, Bourdieu *et al.* ^22^ *report rδ*_*m*_ ∼ 1 − 1.5 nm, *ρ*_*m*_ ∼ 100 − 1000 *µ*m^−2^, and *F*_*m*_ ∼ 0.5 − 0.7 pN, which gives *f* ∼ 10^−6^ Nm^−1^, as well as *ℓ*_*p*_ = 16 *µ*m at room temperature, which gives *κ* ∼ 6 *×* 10^−26^ Nm^2^. Average filament lengths of *ℓ* ∼ 1 − 2 *µ*m seem typical ^43,44^, which yields *β*_max_ ∼ 10^2^. As with microtubules, the length distribution can be broad with some values up to at least 4 *µ*m ^43,44^, which yields *β*_max_ ∼ 10^3^.

Accordingly, our simulations explore 0 ≤ *β* ≤ 10^5^, and it should be noted that some features, most notably self-overlapping filaments, are unique to the upper part of this range (*β* ∼ 10^4^ − 10^5^).

### 2.2 Boundary conditions

The evolution of the filament’s shape and position is completely specified by solving (1)-(2) subject to boundary conditions. The tail is free, i.e., at *s* = *ℓ* the filament is fully unconstrained. The torque and force free conditions thus read

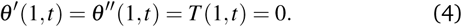

At the head, *s* = 0, we consider three cases: (i) a pinned filament (unable to translate but free to rotate), (ii) a clamped filament (unable to translate or rotate), and (iii) a point viscous drag force and torque on the filament’s head corresponding to a rigidly attached cargo. We first consider case (iii), then discuss cases (i) and (ii) as limit cases of (iii).

The external force **F**_ext_ and the external torque **T**_ext_ exerted at the filament’s head (*s* = 0) by a rigidly attached viscous cargo depend linearly on the cargo’s translational and angular velocities, which in turn depend on the translational and angular velocities of the filament’s head. In the most general case, both **F**_ext_ and **T**_ext_ depend anisotropically on both 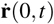 and 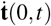. Here, we consider the simpler case of a point viscous load:

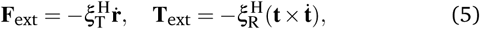

where 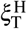 and 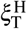 are two scalars representing the head’s effective translational and rotational drag coefficients. Aside from 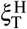 and 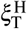, equation (5) does not explicitly involve the geometry of the cargo. Thus by allowing 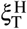 and 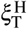 to vary independently we can in principle model a variety of head sizes, head shapes, and friction mechanisms. Even though equation (5) only samples a limited subset of all possible cargo and attachment geometries, we show in section 4 that it is sufficient to reproduce many of the cargo-filament behaviors seen in previous experiments and simulations.

To obtain (5) in terms of *T* and *θ*, we first write 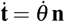 and

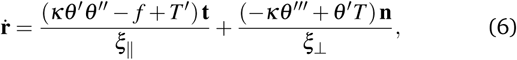

then eliminate the normal force *N* in favor of *θ*. After simplifying and nondimensionalizing (see ESM §II) we obtain

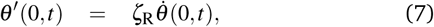

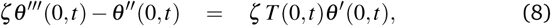

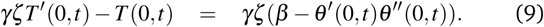

In addition to the dimensionless parameters *γ* and *β* introduced in the bulk Equations (1)-(2), the boundary Equations (7)-(9) feature two new dimensionless parameters: the scaled rotational head drag 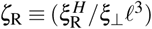 and the scaled translational head drag 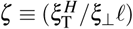. Thus the spatiotemporal evolution of the actively buckling filament is controlled by a total of four dimensionless parameters.

Setting *ζ* = ∞ in Equations (7)-(9) corresponds to holding the filament’s head fixed. (*ζ, ζ*_R_) = (∞, 0) corresponds to a pinned head (no translation, free rotation), while (*ζ, ζ*_R_) = (∞, ∞) corresponds to a clamped head (no translation, no rotation). Thus the head boundary conditions simplify to

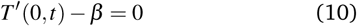

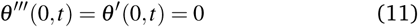

for a pinned head and

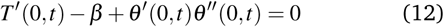

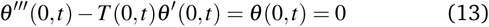

for a clamped head, where without loss of generality we have assumed horizontal clamping (**t**(0) = **e**_*x*_).

## 3 Constrained filaments

Let us first consider constrained filaments (free tail, pinned or clamped head). Similar to classical buckling, we expect the straight configuration to remains table for small force densities (small *β*), then turn unstable to lateral perturbations at some critical value of *β*. Our base state is a straight horizontal filament: *θ* (*s*) = 0 *≡ θ*_0_. From (1)-(2) it follows that *T* (*s*) = *β* (*s*− 1) ≡ *T*_0_(*s*), i.e., the filament is pre-stressed linearly with maximum compression at *s* = 0.

We start with a numerical study of the linear stability of the straight configuration, which predicts the onset of buckling (section 3.1). We follow with an analytical study of the linear stability of a simpler problem with the same stability phenomenology (section 3.2). We then study the numerical solutions of the full nonlinear equations (section 3.3) and relate our problem to other follower-force buckling problems from the literature (section 3.4).

### 3.1 Linear stability

To ascertain any critical points at which the straight pre-stressed filament becomes unstable and to identify the nature of the instability, we write *T* = *T*_0_(*s*) + *εT*_1_(*s, t*) and *θ* = *θ*_0_ + *εe*^*ωt*^ Θ(*s*) where *ε* ≪ 1 is a formal small parameter characterizing the amplitude of the solutions. To first order in *ε*, the equation for Θ decouples from the equation for *T*_1_:

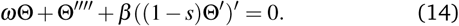

The linearized boundary conditions are given by equation (4) (free tail) and either equation (11) (pinned head) or

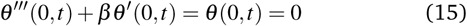

(clamped head). Together, equations (14) and the boundary conditions form an eigenvalue problem. Non-trivial solutions only exist for specific values of the complex eigenvalue *ω*. The stability of the solution is controlled by the sign of the real part of *ω*: stable if it is negative, unstable if it is positive. A nonzero imaginary part indicates an oscillatory solution which may decay or grow depending on the sign of the real part. The shape of the unstable mode is given by the corresponding eigenvector.

We calculate the eigenvalues by discretizing equations (14) and its boundary conditions using a second-order-accurate finite difference method (see ESM §III.A), then track their behavior as a function of the activity parameter *β*. Note that they do not depend on *γ* at all. The first eigenvalue whose real part goes positive signals the onset of instability – a bifurcation point. The manner in which the positive real part arises determines the type of bifurcation.

#### 3.1.1 Pinned filament: Instability to rotational states

In the pinned-free case, the linear stability problem exhibits a unique degeneracy: Θ = Θ_0_≠ 0 is always an eigenvector with eigenvalue *ω* = 0. This Goldstone mode corresponds to a freely rotating straight filament and exists due to the invariance of the elastic energy under rotation about the pinning point.

The blue curves in figures 3(a-b) show the real and imaginary parts of the eigenvalue with the largest real part besides the Gold-stone mode. The straight filament configuration is stable up to *β*_db_ ≈ 30.6, after what it turns unstable as the real part of the eigenvalue becomes positive. The imaginary part remains zero throughout, indicating a divergence bifurcation (DB). At the critical point, both the real and imaginary parts of the growth rate are zero: Re(*ω*) = 0 and Im(*ω*) = 0. As one moves away from the critical point Re(*ω*) ∼ *β* −*β*_db_. Although the eventual shape of the buckled filament is controlled by the nonlinear terms we neglected in this analysis, the zero imaginary part and the presence of the Goldstone mode suggest the filament eventually settles into a steady buckled shape steadily rotating around the pinning point.

**Fig. 3.**
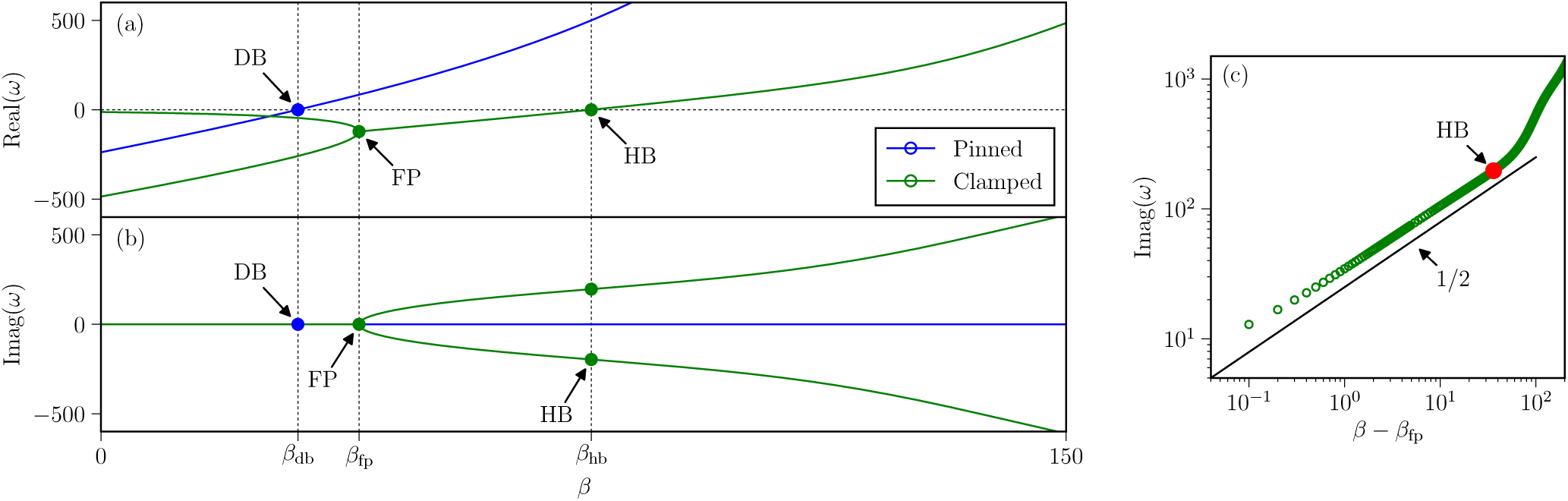
Linear stability of a pinned filament (blue) and a clamped filament (green) as a function of the dimensionless activity *β*. (a) Real part of the most unstable eigenvalue(s). (b) Imaginary part of the most unstable eigenvalue(s). In the pinned case, we discard the zero eigenvalue corresponding to a straight filament. At *β* = *β*_db_ ≈ 30.6, the most unstable eigenvalue acquires a positive real part while maintaining a zero imaginary part, signaling a divergence bifurcation. In the clamped case, the two most unstable eigenvalues start real, distinct, and negative (*β < β*_fp_ ≈ 30.6, overdamped relaxation), then merge their (still negative) real parts and acquire opposite imaginary parts (*β*_fp_ *< β < β*_hb_ ≈ 76.2, damped oscillations), until their real part eventually becomes positive (*β > β*_hb_, unstable oscillations). (c) In the clamped case, the imaginary part scales like (*β* − *β*_fp_)^1*/*2^ near the flutter point consistent with flutter theory ^45^. The scaling remains fairly accurate all the way to the Hopf bifuraction point (red dot).

#### 3.1.2 Clamped filament: Instability to oscillatory states

Clamped-free filaments experience a different type of instability known as a Hopf-Poincaré bifurcation (HB) wherein a pair of complex conjugate eigenvalues crosses the imaginary axis (Re(*ω*) = 0) together. The real and imaginary parts of those two eigenvalues are shown in green in Figures 3(a-b). The instability occurs at *β*_hb_ ≈ 76.2, about two and a half times the load it takes to buckle the pinned filament.

Figures 3(a-b) also show that the pair of complex conjugate eigenvalues that go unstable at *β*_hb_ start off as two distinct, real eigenvalues at small *β*. They only become a complex conjugate pair upon colliding with each other at the flutter point *β*_fp_ ≈ 40.1. Although this happens in the stable regime (Re(*ω*) *<* 0), it signals a change in the filament’s response to fluctuations. Between *β* = 0 and *β* = *β*_fp_, every eigenvalue is real, thus fluctuations relax without oscillating. Between *β* = *β*_fp_ and *β* = *β*_hb_, the slowest relaxation mode (the one whose eigenvalue has the largest real part) exhibits damped oscillations. Finally when *β > β*_hb_ the oscillations become unstable leading to full-blown nonlinear oscillations. This eigenvalue behavior (collision of real eigenvalues turning into a complex conjugate pair) implies the scaling 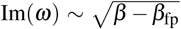 near the flutter point *β*_fp_ ^45^. As seen in Figure 3(c), here this scaling persists until the onset of instability at *β*_hb_ and sets the frequency at the onset of the actual instability: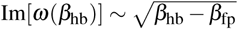.

### 3.2 Minimal models for constrained filaments

Let us now study the linear stability of a simpler problem whose analytical tractability provides a different kind of insight into the nature of the buckling transitions: an elastic filament loaded by a point compressive follower force at the tail. We set *f* = 0 in Equations (1) and (2) and add a tangential compressive force **P** = − *f ℓ***t**(*ℓ, t*) to the boundary condition at *s* = *ℓ*, i.e., we set (in scaled coordinates) *T* (1, *t*) = *β*. The boundary conditions at *s* = 0 remain the same. The crucial change as far as analytical tractabilty is that *T* no longer depends on *s* in the base state. We further eliminate the angle *ψ* in favor of the lateral displacement 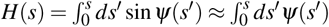. The base state is now given by *H* = 0 and the linearized equation of motion reads

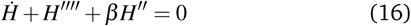

We seek solutions to (16) of the form *H* = exp(*ωt*) *Ĥ* (*s*) and write (without loss of generality) the growth rate as

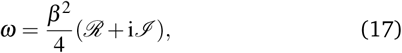

where 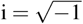. The general solution has the form

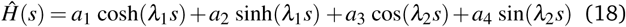

Where

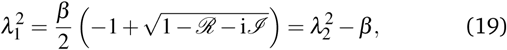

The unknown coefficients in (18) and the values of *λ*_1_ and *λ*_2_ in (18) and (19) are all functions of *β*.

#### 3.2.1 Pinned filament

In the pinned case the boundary conditions are *H*(0) = *H″* (0) = *H″* (1) = *H‴* (1) = 0. Imposing boundary conditions at *s* = 0 on Equations (18) and (19) yields *a*_1_ + *a*_3_ = 0 and 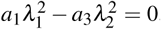. This implies that *a*_1_ = *a*_3_ = 0 since in general 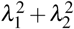 is non-zero. The general form of the solution then reduces to

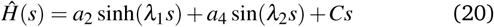

The mode linear in *s* is the Goldstone mode corresponding to a straight rotating filament. It satisfies all the boundary conditions independently.

The solvability condition to keep *a*_2_ and *a*_4_ nontrivial is obtained by applying the boundary conditions at *s* = 1:

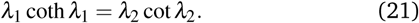

Near a DB point, the growth rate can be estimated by assuming Imag(*ω*) = 0 and Real(*ω*) → 0, or in this case *ℐ* = 0 and *ℛ*≪ *β* ^2^*/*4. Using 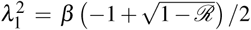,and 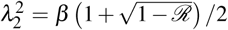 we obtain

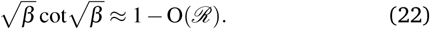

At *ℛ* = 0, equations (22) admits the physical root *β* ≈ 20.19. This is the critical value of *β* at which the straight pinned filament becomes unstable.

#### 3.2.2 Clamped filament

The clamped case was previously studied by De Canio et al. ^25^. Using a method similar to that of section 3.1, they showed that the most unstable eigenvalue exhibits the same qualitative behavior seen in the green curves of Figure 3(a-b). They also derived an equation, similar to equations (22), from which the flutter point and Hopf bifurcation can be obtained without constructing the *ω*(*β*) curves. For the sake of completeness we now rederive this equation in our notations.

The boundary conditions for the clamped case are *H*(0) = *H′* (0) = *H″* (1) = *H‴* (1) = 0. The general solution has the form

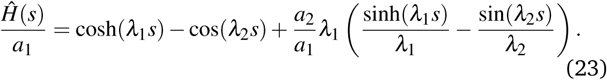

Imposition of boundary conditions at *s* = 1 yields two equations for the two unknowns *a*_1_ and *a*_2_:

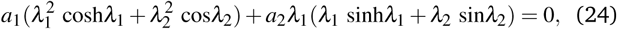

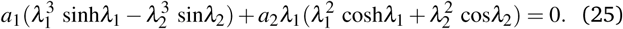

After a bit of algebra, the solvability condition to keep *a*_1_ and *a*_2_ nontrivial can be written as a single complex equation for the three real unknowns *β*, *ℛ, ℐ*:

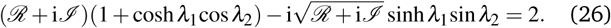

To obtain the flutter point, we set *ℐ* = 10^−6^ (or any value small enough to ensure we are right above the flutter point) and solve for *ℛ* and *β* numerically. For the Hopf bifurcation, we set *ℛ* = 0 and solve for *ℐ* and *β* numerically. We obtain *β*_fp_ = 20.05 and *β*_*hb*_ = 37.70, both consistent with ^25^.

### 3.3 Non-linear solutions and scaling analysis

In the unstable regime, the straight filament yields to buckled solutions whose amplitude is determined by non-linear effects. In order to validate the linear stability analysis and study those non-linear solutions, we integrate Equations (1) and (2) numerically using a semi-implicit finite difference scheme (see ESM §III.B) until a stable solution arises, whether it is a static shape or a periodically evolving one. The initial condition is a slightly perturbed straight filament (*θ* (*s*) = 10^−3^ sin(2*πs*)) that allows instabilities to grow despite the absence of noise in the model.

As expected, at low *β* the initial perturbation relaxes and the filament remains straight. This straight configuration exhibits the same instabilities predicted by the linear stability analysis, at essentially the same force values. The pinned-free filament buckles into a steadily rotating shape at *β*_db_ *≈* 30.6 (linear prediction: *β*_db_ *≈* 30.6). The clamped-free filament buckles into a periodic beating pattern at *β*_db_ *≈* 76.3 (linear prediction: *β*_db_ *≈* 76.2). Figures 4(a-b) show typical stable filament shapes from a little above the onset of each instability up to very large *β*. In the pinned-free case (figure 4(a)), the shape is static in the co-rotating frame, thus a single snapshot suffices. For ease of comparison every filament is shown with its head pointing to the left. In the clamped-free case (figure 4(b)), multiple snapshots are needed to visualize the beating pattern. At β = 75 and 150 we show three such snapshots. At *β* = 105 we show a single snapshot illustrating the filament intersecting itself. In both pinned and clamped filaments, the shapes show increased curvature as *β* increases, confirmed by the increasing bending energy shown in Figure 4(c), until the filament eventually overlaps itself (rightmost shapes in Figures 4(a-b)). Note that this model does not prevent self-overlap at all. In section 5 we discuss a model with short-range repulsion that is better suited for systems in which self-crossing is effectively forbidden.

**Fig. 4.**
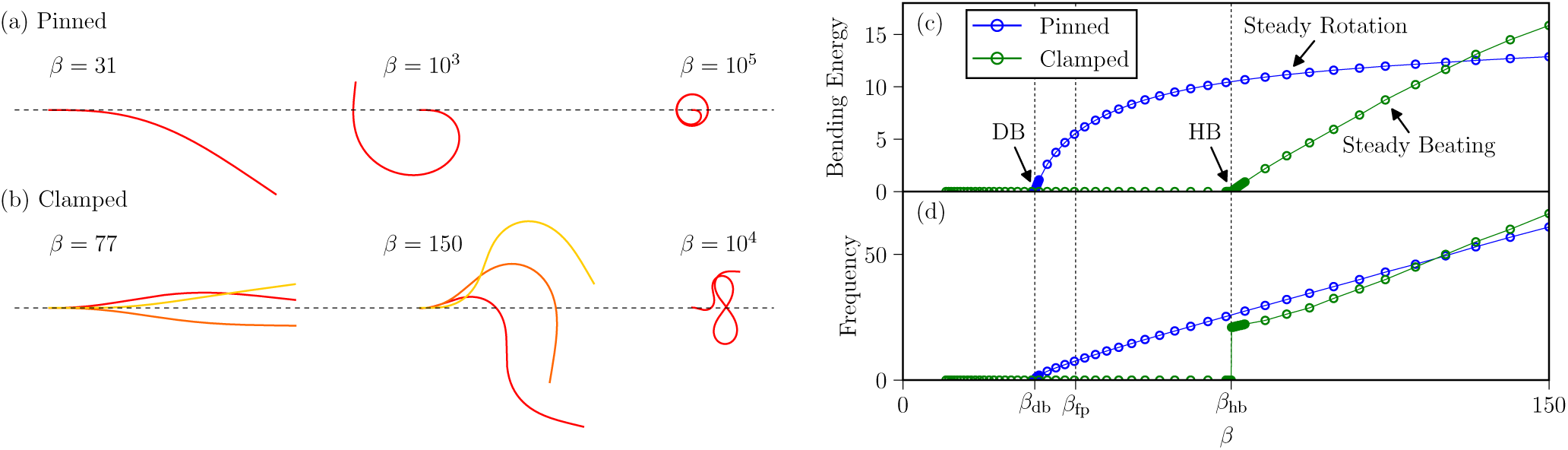
Features of the fully non-linear, steady state solutions for *γ* = 2. (a,b) Typical filament shapes for (a) a pinned filament and (b) a clamped filament for activity parameters *β* varying from close to the critical value to values at which self-crossing occurs. In the pinned case, the steady-state behavior is solid body rotation with constant-sign curvature. At very large *β* the filament rolls into a self-overlapping coil. In the clamped case, the steady-state behavior is periodic beating. Each color (red, orange, yellow) is a different configuration (shape) comprising the beating pattern. Again, increasing the driving force eventually leads to self-crossing. (c) Mean bending energy as a function of *β*. (d) Rotation (pinned case) or beating (clamped case) frequency as a function of *β*

A number of large-*β* properties can be understood through scaling arguments. Balancing the active force density *f* and the tangential bending force density ***κ*** *θ* ′ *θ* ″ ∼ ***κ*** *θ* ^2^/ *λ*^3^ in a filament with typical curvature radius *λ* yields *λ* =(*κ*/ *f*) ^1/3^ = *𝓁*/ *β* ^1/3^. This is the same *λ* we introduced below equations (3) to discuss the meaning of *β*. When *λ* is larger than the filament’s length *𝓁, β* is small and the filament remains straight. When λ is sufficiently shorter than the filament’s length *𝓁, β* is large, the filament buckles, and we expect its curvature radius to be of order *λ*. We can then use *λ* to estimate the bending energy: 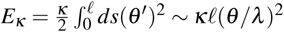 At large *β* we expect *θ* ∼ 1 thus *E*_*κ*_ ∼ *κ 𝓁 λ* ^2^= (*κ*/*𝓁*) *β*^2/3^

The scaling exponents we observe in Figure 5(a-b) are a little smaller than 2*/*3. Looking at the spatial variations of the curvature along the filament (*θ*^*t*^(*s*), shown in Figure 5(c-d)), it is clear that even at the largest *β* we simulated (*β* = 10^5^), a good 15% of the filament experiences significant boundary effects. Thus we think the exponent discrepancy is due to boundary effects and expect the 2*/*3 exponent to be observable at yet higher *β*.

**Fig. 5.**
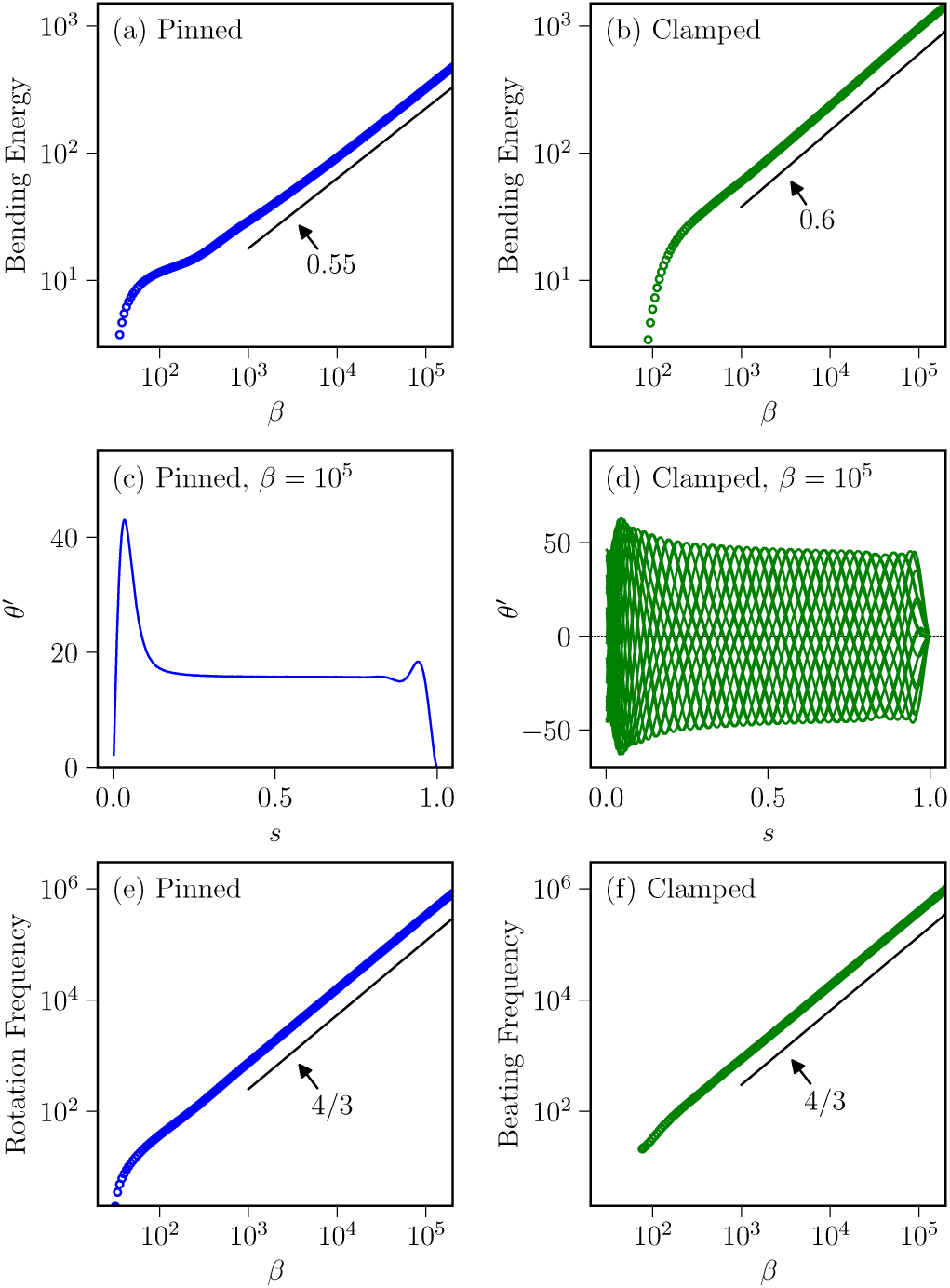
Features of the fully non-linear, steady-state solutions for a pinned filament (blue) and a clamped filament (green) for *γ* = 2. (a-b) Mean bending energy as a function of the active force density *β*. The scaling exponents are slightly smaller than the predicted 2*/*3. (c-d) Curvature *θ* ^*′*^ as a function of the arclength *s* along the filament for the largest force density we simulated (*β* = 10^5^). In the clamped case we show 20 different times during the beating pattern to visualize the envelope of the traveling curvature wave. In both cases boundary effects are still significant, which may explain the discrepancy with the predicted scaling exponents in (a-b). (e-f) Rotation (pinned case) and beating (clamped case) frequency as a function of *β*. In both cases the predicted 4*/*3 scaling exponent is accurate.

To estimate the rotation frequency of pinned filaments at high *β*, we start from the coil shape shown in Figure 4(a). Assuming the coil has radius *λ*, its linear (tangential) speed is *v* ∼ (*ℓ f*)*/*(*ℓ ξ* _‖_) and its angular frequency is *ω* ∼ *v/*λ** ∼ *κ/*(*ξ _‖_ ℓ*^4^) *β* ^4/3^.

To estimate the beating frequency of clamped filaments, we note that for the bending energy to come back to the same value at the end of every beat, the energy *E*_*f*_ supplied by the active force over a full beat must equate the energy *E*_*ξ*_ dissipated by viscous drag over the same period. The former is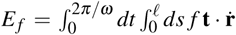 Assuming the total distance traveled by a point of the filament over a full beat is of order *λ*, we can write 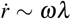 then *E*_*f*_ ∼ (1*/ω*)(*ℓ*)(*f*)(*ω*λ**) = (*κ/ℓ*)*β* ^2*/*3^. The latter is 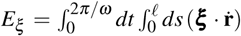 where *ξ* = *ξ* _*‖*_**tt** + *ξ*_*⊥*_(1 − **t t**) is the filament’s drag tensor. Using the same estimates we get *E*_*ξ*_ ∼ (1*/ω*)(*ℓ*)(*ξ*)(*ω*λ**)^2^ = *ℓ*^3^*ξωβ*^−2*/*3^ where *ξ* is a combination of *ξ _‖_* and *ξ*_*⊥*_. Finally we equate the two energies to get *ω* ∼ *κ/*(*ξℓ*^4^) *β* ^4*/*3^.

As shown in Figure 5(e-f), the rotation frequency of pinned filaments and the beating frequency of clamped filaments do both grow like *β* ^4*/*3^ as early as *β* ∼ 10^3^.

### 3.4. Related constrained follower-force problems

The linear stability problem of section 3.1 was analyzed by Sekimoto *et al.* ^24^ *who found *β**_db_ *≈* 30.6 and *β*_hb_ *≈* 75.5. The small discrepancy in *β*_hb_ may be due to a different discretization method. De Canio *et al.* ^25^ studied the linear stability and nonlinear dynamics of the clamped point follower load model of section 3.2.2 and found the same qualitative behavior we see in the clamped distributed follower force case. Ling *et al.* ^26^ studied the linear stability and nonlinear dynamics of the 3D clamped case and found two distinct types of beating, a planar one similar to the one we observe and a helical one, each having its own instability threshold and a range of *β* in which it dominates. Fatehiboroujeni *et al.* ^46^ studied the double-clamped 3D case, in which both ends of the filament are clamped and the base state is pre-buckled. Chellakot *et al.* ^27^ studied a slightly different problem with noisy follower forces using a bead-spring model (discussed further in section 5) and found *β*_db_ *≈* 36 and *β*_hb_ *≈* 78 in the weak (but finite) follower force noise regime. Reference ^27^ also discusses the *β* ^4*/*3^ growth of the frequency in both pinned and clamped filaments, however they find it no longer holds at very large *β*. We suspect this restriction comes from either the discrete nature of their bead-spring model, the presence of steric repulsion, or both.

Most of the literature on buckling under follower forces, however, comes from engineering ^39^. In this context, an elastic filament under compressive follower load is known as a Leipholz column if the load is distributed along the filament as in section 3.1 or a Beck column if the load is concentrated at one end as in section 3.2. The latter may arise, for example, when a jet engine is attached at the end of an elastic column. A key difference with the present paper is that those studies are typically concerned with moderate or weak damping, whereas we’re interested in the overdamped limit when inertia is negligible.

To better understand the relationship between the two situations, it is useful to consider the undamped version of our problem, i.e., replace the anisotropic friction force [*ξ _‖_***tt** + *ξ*_*⊥*_(𝟙 − **tt**)]. *∂*_*t*_ **r** with an inertial term 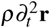 where *ρ* is the mass density per unit length of the filament. Up to some redefinitions of the dimensionless parameters, this is equivalent to changing 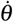 to 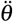 in equations (2) and *ω* to *ω*^2^ in equations (14) ^46^. The free, pinned, and clamped boundary conditions are unchanged. In other words, the inertial-case growth rate is the square root of the overdamped-case growth rate with the same dimensionless parameter values. The same reasoning applies when the follower load is concentrated at the end of the filament.

In the pinned-free case, the eigenvalues before the divergence bifurcation point (*β* < *β*_db_) are all real and negative, therefore the corresponding inertial growth rates are purely imaginary, corresponding to constant-amplitude oscillations of the inertial filament. At the bifurcation (*β* = *β*_db_), one eigenvalue turns positive, thus one of its square roots turns real positive. Therefore, the inertial system goes unstable at the same dimensionless force *β* as the overdamped one, and the unstable growth rate is real in both cases.

In the clamped free case, the eigenvalues before the flutter point (*β* < *β*_fp_) are all real and negative. This, again, suggests constant-amplitude oscillations of the inertial filament. At the flutter point, two eigenvalues acquire opposite nonzero imaginary parts while their real part remains negative. This results in a pair of complex conjugate square roots with positive real part. Both the overdamped and the inertial system experience a transition at *β*_fp_, however the overdamped system merely transitions from overdamped stability to oscillatory stability, whereas the inertial system goes unstable altogether. The change of sign of the real part of this pair of eigenvalues at *β* = *β*_hb_ then has no major significance to the inertial system.

In summary, the effect of strong damping depends strongly on the boundary conditions. For pinned-free filaments the change is essentially limited to pertubations in the stable region relaxing rather than oscillating. For clamped-free filaments, on the other hand, damping impacts both the location and the nature of the transition.

## 4 Filaments moving attached cargo

Let us now consider filaments whose head, rather than being held fixed (pinned or clamped), is attached to a viscous load, for example an active filament transporting a cargo. The effect of the head is captured by the scaled translational and rotational drag paramaters *ζ* and *ζ*_R_ from Equations (7)-(9). With the activity parameter *β* and the viscosity contrast *γ*, this makes four dimensionless parameters.

As before, the stable configuration under weak loading (small *β*) is straight, however it now translates at constant speed along its own length:

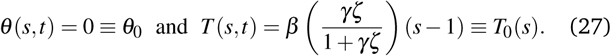

Substituting (27) in (6) yields the velocity of the head, which is also that of the entire filament:

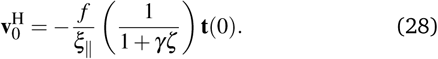

Equations (27)-(28) define the base state whose instabilities we now study.

### 4.1 Methods

Following section 3, we study both the linear stability of the base state and the long-time solutions of the full nonlinear equations.

To analyze the linear stability of the base state, we write *θ* (*s, t*) = *θ*_0_ + *ε*Θ(*s, t*) with *ε* ≪ 1, substitute Equations (27)-(28) into Equations (1)-(2) and (7)-(9), and expand to *O*(*ε*) to get

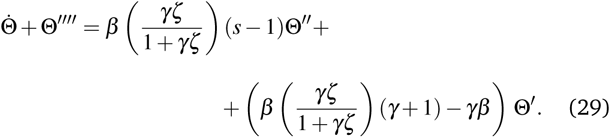

and

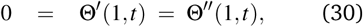

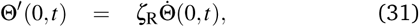

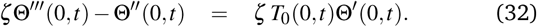

We then discretize those linearized equations as in section 3.1, compute the eigenvalues, and extract the critical value(s) corresponding to the onset of instability.

To obtain the full nonlinear solutions we discretize Equations (1)-(2) and (7)-(9) as in section 3.3.

#### 4.1.1 Results

Figures 6(a-c) show the regions of (in)stability in the (*β*, *ζ*_R_) plane for *γ* = 2 and three different values of *ζ* (from left to right: 0.2, 1, 5). Each colored dot corresponds to a nonlinear simulation: light gray for filaments that remain straight, red for filaments that buckle into a rotating state, blue for filament that buckle into a beating state. Rotating filaments look similar to the ones in Figure 4(a) except their head now follows a circular orbit (see ESM Movie 1). Beating filaments exhibit periodic buckling similar that of Figure 4(b) except their head now moves along a wavy line (see ESM Movie 2).

**Fig. 6.**
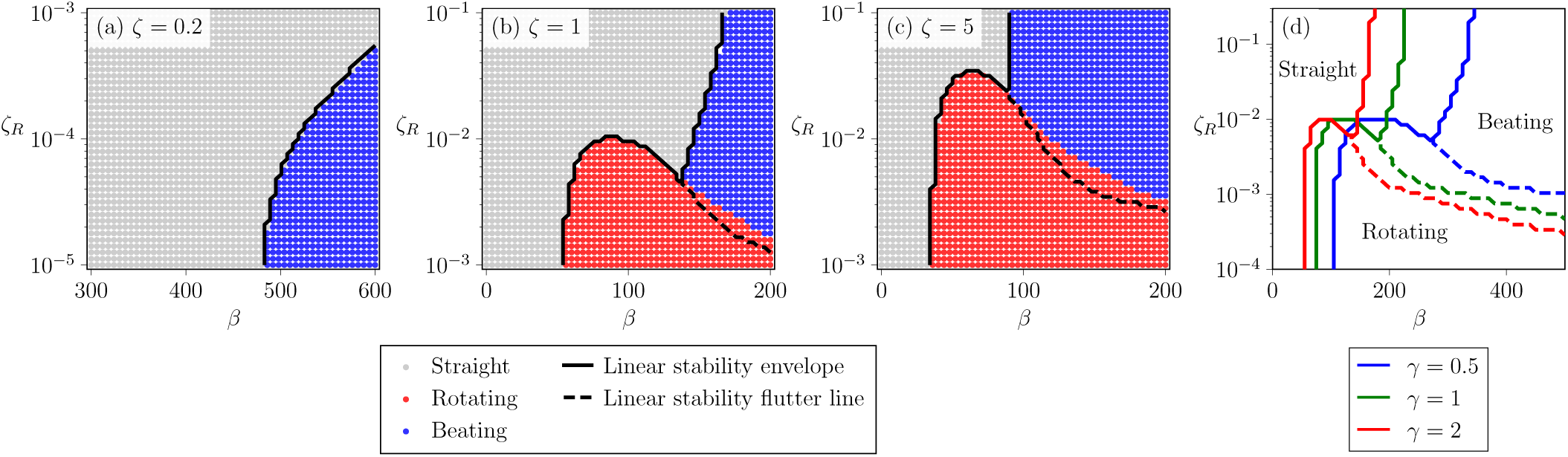
(a-c) Instabilities of a filament attached to a viscous head as a function of the active force density *β* and the head’s rotational drag *ζ*_R_ for *γ* = 2 and three values of the head’s translational drag *ζ* (left: 0.2, center: 1, right: 5). Each colored dot represents a nonlinear simulation. The color codes for the long-time behavior: light grey for straight translation, red for buckled rotation, blue for buckled beating. The black lines represent the boundaries of the linear stability regions. The solid lines separate the stable region from the two unstable regions. The dashed lines separate the two unstable regions; the imaginary part of the most unstable eigenvalue is zero below the line, nonzero above it. (d) Linear stability boundaries between the three states of a filament-cargo assembly (straight, rotating, beating) in the (*ζ*_*R*_, *β*) plane for *ζ* = 1 three values of the viscosity contrast *γ*. Increasing the filament’s normal to tangential drag coefficient ratio *γ* shifts the phase boundaries towards lower values of the active force *β*.

The linear stability analysis is performed on a similar grid of (*β, ζ*_*R*_) values, however for readability only the boundaries of the (in)stability regions are shown. The solid black curve corresponds to the appearance of an eigenvalue with a positive real part, i.e., the straight/buckled boundary. The dashed black curve corresponds to the most unstable eigenvalue acquiring a nonzero imaginary part, i.e., the rotating/beating boundary.

Many of the features seen in Figures 6(a-c) can be related to the results of section 3. First, *β* = 0 always yields a straight filament. Second, following the bottom edge of Figure 6(c) corresponds to varying *β* while holding *ζ* and *ζ*_R_ fixed with *ζ* large and *ζ*_R_ small.

This may be thought of as the finite-*ζ*, finite-*ζ*_R_ version of the pinned boundary condition (*ζ* = ∞, *ζ*_R_ = 0). Accordingly, both situations yield a transition from a straight filament to a rotating one as *β* is increased. Likewise, the finite (*ζ, ζ*_R_) counterpart of the transition from a straight clamped filament to a beating one can be observed along the top edge of Figure 6(c). The overall shape of Figures 6(b) and 6(c) follows from bridging those three limits: straight states on the left (low *β*), rotating states at the bottom right (large *β*, low *ζ*_R_), and beating states at the top right (large *β*, large *ζ*_R_).

Still, there are some unexpected features. First, we could not identify a rotating region at *ζ* = 0.2, suggesting it may disappear entirely at low *ζ*. Second, the straight state is re-entrant. For example, at fixed *ζ* = 1 and *ζ*_R_ = 0.007, a straight filament remains straight if *β* is small, buckles into a rotating shape if *β* is a little larger, remains straight if *β* is yet larger, and buckles into a beating shape if *β* is even larger. In other words, an unstable straight filament can sometimes be re-stabilized by further increasing the active buckling load (*β*). Third, near the transition between rotating and beating we observe mixed states wherein the filament switches periodically between rotating and beating. ESM Movie 3 illustrates this behavior, however note that trajectories in this region are quite sensitive to the amount of time spent in each state and the timing of the switches.

It is also worth noting that the linear stability analysis performs well at the straight/buckled transition, but not so well at the rotating/beating transition. This is because the latter involves two highly curved states, neither of which is well described by a first order expansion around the straight state.

Finally, Figure 6(d) illustrates the effect of the viscosity contrast *γ*. It shows the linear stability boundaries in the (*β, ζ*_*R*_) plane for *ζ* = 1 and three different values of *γ*: 0.5 (blue), 1 (green), and 2 (red, same as the black lines in Figure 6(b)). There are no qualitative changes, only an overall shift of the phase boundaries towards weaker *β* values as *γ* increases with the straight/beating line shifting more than the straight/rotating line.

## 5 Robustness to noise and steric repulsion

The nonlinear simulations of sections 3-4 do not include any noise. Neither do they include steric repulsion preventing the filament from overlapping itself. Whether the latter is relevant in an experiment depends on the details of the quasi-two-dimensional confinement, however noise is always present at the micro-scale. We now show that our results are robust, at least qualitatively, to both. Rather than adding those effects to our continuum model, we switch to a bead-spring model inspired by previous work by some of us ^27^.

### 5.1 Model

The model is a simplified version of those introduced by Chelakkot *et al.* ^27^ *and Isele-Holder et al.* ^28^. *It consists* of a chain of elastically connected, self-propelled, polar colloidal spheres that moves and deforms in two dimensions. The filament has *N* spheres, each of diameter *σ*, located at coordinates **r**_*i*_ (*i* = 1, *.., N*). Adjacent beads are connected by linear springs with equilibrium length *σ* and stiffness *κ*_E_. The corresponding potential is

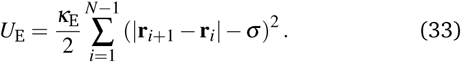

*κ*_E_ is chosen large enough to keep the bonds’ lengths approximately equal to *σ*, thus mimicking inextensibility. The chain’s length is then *ℓ ≈ Nσ*.

Resistance to bending is implemented via a three-body elastic potential with bending rigidity *κ*.

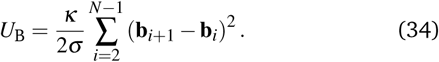

where **b**_*i*_ = (**r**_*i*−1_ − **r**_*i*_)*/*|**r**_*i*_ − **r**_*i*−1_| is the unit bond vector, which always points towards the next bead in the direction of the head bead (bead 1). In the continuous limit (*σ →* 0, *N →* ∞, *Nσ →*constant), **b**_*i*_ identifies with the tangent vector **t** of the continuous model at arclength *s* = *iσ*, thus (**b**_*i*+1_ − **b**_*i*_)*/σ ≈ d***t***/ds* and *U*_B_ *≈* (*κ/*2) *ds*(*d***t***/ds*) = (*κ/*2) *ds*(*d*θ*/ds*)^2^. In other words, the definition (34) ensure *κ* has the same meaning as in the continuous model.

The active follower force pushes each bead with magnitude *F* along its bond vector: **F**_*i*_ = *F***b**_*i*_. In the continuous limit, this yields a uniform active force per unit length *f* = *F/σ* which identifies with the active force density used in the continuous model.

Steric repulsion takes the form of a short-ranged pairwise repulsive WCA (Weeks-Chandler-Anderson) interaction between beads. The potential is

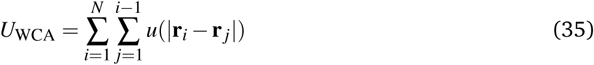

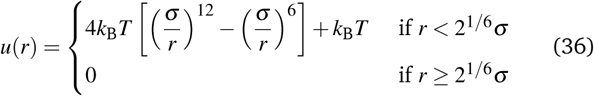

where *k*_B_ is Boltzmann’s constant and *T* is the temperature.

The positions of the beads evolve according to over-damped Brownian dynamics:

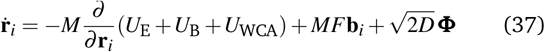

where *M* is the viscous mobility, *D* is the diffusion constant, which obeys Stokes-Einstein’s equation *D* = *Mk*_B_*T*, and Φ is a zeromean, unit-variance Gaussian white noise force.

The cargo (viscous load) consists of a hexagon made of seven beads rigidly attached to each other and to the filament’s first bead (*i* = 1, corresponding to *s* = 0 in the continuous model) as shown in Figure 7. The cargo beads experience noise, drag, and WCA interactions like the filament’s beads, however they do not experience any active follower, spring, or bending spring force.

**Fig. 7.**
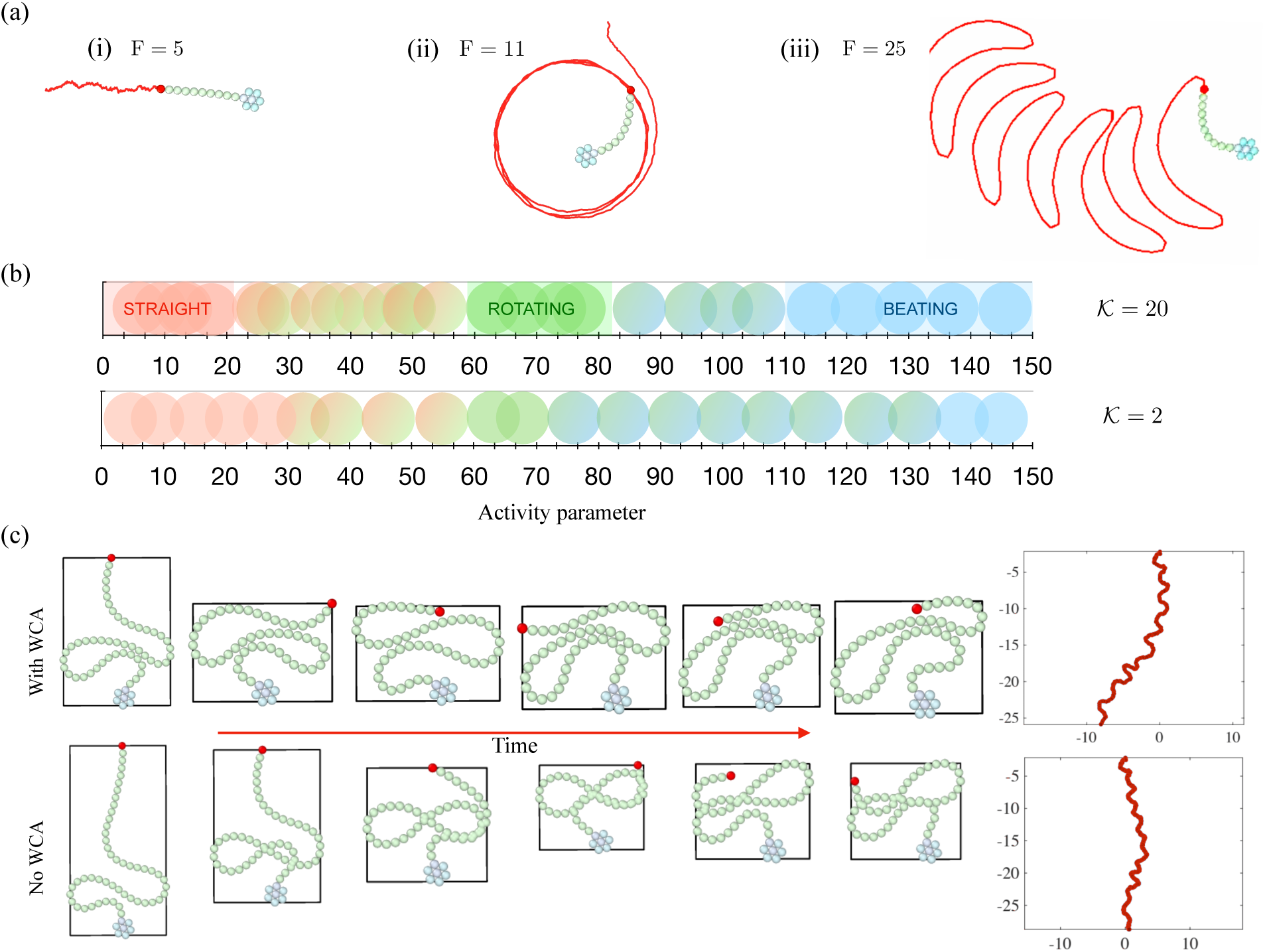
(a) States of a noisy swimmer comprised of a 10-bead filament pushing a 7-bead hexagonal cargo: straight at low force density *F* (left), rotating at intermediate *F* (middle), beating at high *F* (right). The red bead is the tail; the red curve is its trajectory. (b) Phase diagram of the same swimmer as a function of *F* with weak (top, *𝒦* = 20) and moderate (bottom, *𝒦* = 2) noise. Red, green and blue represent straight, rotating, and beating states respectively. Mixed colors represent transition states. (c) Beating pattern of a 50-bead filament pushing a 7-bead hexagonal head at *F* = 40 with (top) and without (bottom) WCA repulsion. The red curves on the right represent the planar trajectory of the tail bead. Length scales are measured in units of *σ*.

Equations (34)-(38) are rendered dimensionless by scaling lengths by *σ*, times by *σ* ^2^*/D*, and energies by *k*_B_*T*. In the simulations this is done by setting *σ* = 1, *M* = 1, and *k*_B_*T* = 1. We are then left with three dimensionless parameters: the dimensionless stretching stiffness *κ*_E_*/*(*k*_B_*T*), the dimensionless bending rigidity *κ/*(*σk*_B_*T*), and the dimensionless active force *σ F/*(*k*_B_*T*). The exact value of the former is not very important as long as it is large enough to keep length fluctuations small and the chain approximately inextensible. We set *κ*_E_*/*(*k*_B_*T*) = 20000. The other two parameters, while convenient to run the simulations, are not best suited to analyze the results and compare with the continuous model. Thus we define the dimensionless activity parameter *β* = *Fσ* ^2^*N*^3^*/κ*, which identifies with our previous *β* in the continuous limit, and the (new) dimensionless bending rigidity *𝒦* = *κ/*(*Nσk*_B_*T*). The latter is the ratio of the thermal persistence length *ℓ*_*p*_ *= κ/*(*k*_B_*T*) to the chain’s length *ℓ* = *Nσ*. If *𝒦* ≫ 1 (large bending rigidity, short chain, or weak noise), the noise alone does not preclude a straight chain. Conversely, if *𝒦* ≪ 1 (small bending rigidity, long chain, or strong noise) thermal noise alone can destroy the straight configuration. Since the relative noise strength is set by 1*/ 𝒦*, one may explore the role of noise by varying any of *κ, k*_B_*T*, or *ℓ*. Isele-Holder *et al.* ^28^ *fixed κ* and *L* and varied *k*_B_*T*. Chelakkot *et al.* ^27^ *fixed σ* and *k*_B_*T*. Here we follow the latter.

The simulations start with a straight filament then use the Euler-Maruyama scheme to integrate the equations of motion until a stable motion pattern is established. Each set of parameters is simulated for ∼ 10 distinct realizations of the noise to ascertain the validity of the observed pattern.

## 5.2 Results

We start with a ten-bead filament (*N* = 10). At both *𝒦* = 20 (low noise) and *𝒦* = 2 (moderate noise), the filament goes through all three swimming states as *F* increases from 0: straight, then rotating, then beating. Figure 7(a) and ESM Movie 4 show typical trajectories in each state. Figure 7(b) shows the phase diagram along the *F* axis. The same sequence of states could be observed by following a horizontal line in the lower region of Figure 6(b) so as to cross the dashed line. The intermittent switching between rotating and beating we observed across that dashed line in section 3 is still present in the noisy simulations. In fact, a similar behavior is now present at the straight-rotating transition, except it is now driven by noise, thus random. Overall, the effect of noise on the phase diagram is to broaden the transitions. Unsurprisingly, this broadening is stronger at *𝒦* = 2 than at *𝒦* = 20.

Although we did not explore the (*ζ, ζ*_*R*_, *β*) phase diagram as systematically as we did in Figure 6, the results above suggest the phase diagram is at least somewhat robust to noise, and we expect no changes to the overall shape of Figure 6 in presence of weak to moderate noise.

The ten-bead simulations above include both noise and steric repulsion, however the latter is irrelevant as the filament is too short and the active force values we simulated far too weak for the filament to fold back onto itself. To explore steric effects, it is convenient to switch to a 50-bead filament with the same 7-bead hexagonal head and *F* = 40, which yields large-amplitude beating. Figure 7(c) shows the filament’s shape at various times of its beating pattern and the trajectory of it tail (rightmost panel). In the bottom row, the WCA repulsion is turned off. Predictably, steric repulsion increases the space occupied by the filament. This in turn increases the beating wavelength, thus the curvature length scale. However, it does not seem to disrupt the beating itself. Similar observations can be made about highly curved rotating filaments: steric repulsion increases the space occupied by the filament and its radius of curvature, however it does not prevent the filament from rotating.

## 6 Discussion

In summary, the continuous model of section 2 and the stability results encapsulated in Figure 6 successfully capture the qualitative phase diagram and the essential dynamical features of 2D elastic filaments driven by follower forces as have been observed in more detailed simulations and in experiments.

Much of the dynamics seen in ^27^, at least in the weak motor noise regime, can be understood in terms of the linear stability analysis of section 3.1 and the scaling analysis of section 3.3. The qualitative features of the linear stability analysis are reproduced by the simplified models of section 3.2, whose analytical tractability can provide further insight into the emergence of the divergence bifurcation seen in pinned filaments and the flutter point and Hopf bifurcation seen in clamped filaments.

Both pinned rotating (figure 1) and clamped beating ^47^ states have also been observed in gliding motility assay experiments, although the transient nature of some anchoring sites can complicate matters. In Figure 1, for example, the microtubule first encounters a pinning defect and starts to form a rotating coil, but in the third panel the head has freed itself from the pinning defect and resumed its straight motion.

For filaments pushing a cargo, Isele-Holder et al. ^28^ found numerical evidence of the straight, rotating, beating, and rotating/beating intermittent states by varying the size and shape of the cargo attached to a bead-spring filament. Rather than focusing on any particular head shape or size, we note that the effect of any viscous-drag-generating rigidly-attached head can be captured by a finite set of drag parameters. Although a full exploration of the cargo parameter space is beyond the scope of this paper, our systematic exploration of the role of two such parameters (ζ and ζ_R_), along with the two dimensionless parameters characterizing the filament itself (*β* and γ), proves sufficient to observe many of the dynamical states seen in previous simulations and experiments. It also provides a solid starting point for a future, more complete exploration of the cargo parameter space. Perhaps most importantly, this approach could prove useful to design filament-cargo assemblies exhibiting a specific behavior by breaking down the problem into two pieces: first, identify a region in cargo parameter space that produces the desired behavior; then, look for a cargo shape and size whose drag parameters are in said region.

We conclude with some possible extensions of this work. First, one may generalize our treatment of filament-cargo attachment by varying the entire set of drag parameters needed to describe any rigidly-attached viscous head as discussed in section 2.2. Second, the 2D constraint may be relaxed to allow motion in 3D. Ling et al. ^26^ found two distinct beating instabilities, one of them helical, in the constrained problem, however we are not aware of any such study for filament-cargo assemblies. Third, long-range hydrodynamic interactions may be included, e.g., by assuming the head is spherical and adding stresslets distributed along the filament ^48–50^. This may become important for large cargos, or at high filament density when filament-filament interactions are frequent. Fourth, one may check the robustness of our cargo results to noise in follower force due, e.g., to fluctations in the motor density. This would also extend ^27^, which includes this noise but only considers constrained filaments. Finally, arc-length continuation techniques may be used to identify unstable dynamical patterns or isolated stable branches that are not easily accessible by integrating time starting from a straight filament.

## Acknowledgements

AG and PS would like to thank S. Koch for sharing the gliding assay image series. AG acknowledges support from UC Merced startup funds. PS would also like to acknowledge funding from a L’Oréal-UNESCO Women in Science UK fellowship. Initial work on this paper was started by AG, PS and TS at MPIDS Goettingen.

## Supporting information

Electronic Supplementary Information

ESM Movie 1

ESM movie 2

ESM Movie 3

ESM Movie 4

## Author Contributions

AG and PS conceived the initial study with TMS at MPIDS Goettingen. AG designed the theoretical model. YF, AG and PS performed the linear stability analysis. YF and AG developed code to obtain and analyze the non-linear solutions. AG and RC wrote the Brownian dynamics code and AG analyzed its results. All authors approve the manuscript as written.

## Data availability

The authors declare no competing financial interests. Correspondence should be addressed to AG. All relevant data and simulation details are available from YF and AG upon request.

