## Supplementary material for "Instabilities and Spatiotemporal Dynamics of Active Elastic Filaments": Electronic Supplementary Information

### ELECTRONIC SUPPLEMENTARY MATERIAL

Yaouen Fily[a†], Priya Subramanian[b†], Tobias M. Schneider[c], Raghunath Chelakkot[d], Arvind Gopinath[e*]
[a] *Wilkes Honors College, Florida Atlantic University, Florida, USA*
[b] *Mathematical Institute, University of Oxford, Oxford OX2 6GG, United Kingdom*
[c] *Emergent Complexity in Physical Systems Laboratory (ECPS), Ecole Polytechnique Federale de Lausanne, CH 1015 Lausanne, Switzerland.*
[d] *Department of Physics, IIT Bombay, Mumbai, India.*
[e] *Department of Bioengineering, University of California, Merced, CA, USA.*

[†] *Equal contribution*
[*] **

### I. GOVERNING EQUATIONS

In this section, we derive the dimensional form of the equations governing the dynamics of the animated, active filament of length $\ell$ and bending modulus $\kappa$ using force and torque balances.

We work in the local orthonormal coordinate system defined at each arclength $s$ by the local tangent $\mathbf{t}(s,t)$ and the local normal $\mathbf{n}(s,t)$ to the filament. We start by decomposing the internal force resultant $\mathbf{F}(s,t)$ (area integral of the stress tensor) at the material point located at arc-length position $s$,

$$\mathbf{F} = T\mathbf{t} + N\mathbf{n}, \tag{1}$$

into its tangential $T(s,t)$ and normal $N(s,t)$ components as shown in Figure 2 of the main text. $\mathbf{F}(s)$ is the force exerted by the tail side of the filament (between arclength $s$ and 1) onto the head side of the filament (between arclengths 0 and $s$), and $T$ is the filament's tension. Similarly we define the bending moment $\mathbf{M}(s)$ as the torque exerted by the tailside of the filament onto its headside at $s$. For a slender elastic beam it is given by $\mathbf{M} = \kappa\,\theta'\,\mathbf{t}\times\mathbf{n}$.

Next we write force balance for the differential element $ds$ located between $s$ and $s+ds$:

$$-\mathbf{F}(s) + \mathbf{F}(s+ds) - f\mathbf{t}(s)\,ds + \mathbf{f}_v(s)\,ds = \mathbf{0}. \tag{2}$$

The active force density always acts along the local tangent vector with a consistent direction (towards $s=0$) and constant magnitude $f$. The friction force per unit length is an anisotropic linear function of the local filament velocity:

$$\mathbf{f}_v = -(\xi_{\|}u_{\|}\mathbf{t} + \xi_{\perp}u_{\perp}\mathbf{n}) \tag{3}$$

where $\xi_{\|}$ and $\xi_{\perp}$ are the effective viscous resistances per unit length for motion of the filament along its local tangent and normal vectors, respectively, and $u_{\|}$ and $u_{\perp}$ are the components of the local velocity of the centerline in the local $(\mathbf{t},\mathbf{n})$ frame. Invoking the slender body assumption allows us to express the arc-length variations of $\mathbf{F}$ in terms of the force components and the shape of the filament parametrized here by the angle $\theta$ between the local tangent vector and the positive x axis:

$$\mathbf{F}' = (T' - N\theta')\,\mathbf{t} + (N' + T\theta')\,\mathbf{n}. \tag{4}$$

Combining (2)-(4) we obtain equations relating the shape variable $\theta$ and the state variables $N$ and $T$ to the tangential and normal components of the velocity of the centerline:

$$(N' + T\theta') - \xi_{\perp}u_{\perp} = 0, \tag{5}$$

$$(T' - N\theta') - f - \xi_{\|}u_{\|} = 0. \tag{6}$$

Next we write torque balance for the same differential element $ds$:

$$-\mathbf{M}(s) - \mathbf{M}(s+ds) + (ds\,\mathbf{t})\times(N\mathbf{n} + T\mathbf{t}) = 0\,. \tag{7}$$

This yields $N = -M' = -\kappa\theta''$, which we use to eliminate $N$ from (5)-(6):

$$-\kappa\theta''' + T\theta' = \xi_{\perp}u_{\perp}, \tag{8}$$

$$T' + \kappa\theta''\theta' - f = \xi_{\|}u_{\|}. \tag{9}$$

To close this set of equations, we use the inextensibility condition $\mathbf{r}' = \mathbf{t}$, which yields

$$\dot{\mathbf{t}} = (u_{\|}\mathbf{t} + u_{\perp}\mathbf{n})' = (u_{\|}' - \theta' u_{\perp})\,\mathbf{t} + (u_{\|}\theta' + u_{\perp}')\,\mathbf{n}, \tag{10}$$

Combining with $\dot{\mathbf{t}} = \dot{\theta}\mathbf{n}$ provides a relationship between the velocity components and the shape:

$$\dot{\theta} - (u_{\perp}' + u_{\|}\theta') = 0, \quad \text{and} \quad u_{\|}' - u_{\perp}\theta' = 0. \tag{11}$$

Finally we use (11) to eliminate the filament's velocity from (8)-(9) and obtain

$$T'' = -\kappa(\theta''\theta')' + f' + \frac{\xi_\parallel}{\xi_\perp}\,\theta'\,(-\kappa\theta''' + T\theta')\,, \tag{12}$$

$$\xi_\perp\dot{\theta} = -\kappa\theta'''' + (T\theta')' + \frac{\xi_\perp}{\xi_\parallel}\,\theta'\,(T' + \kappa\theta''\theta' - f), \tag{13}$$

$$\dot{\mathbf{r}} = \frac{(\kappa\theta'\theta'' - f + T')\,\mathbf{t}}{\xi_\parallel} + \frac{(-\kappa\theta''' + \theta' T)\,\mathbf{n}}{\xi_\perp}. \tag{14}$$

#### II. BOUNDARY CONDITION FOR FILAMENT-CARGO ASSEMBLIES

Here we assume a point viscous load is attached at $s = 0$ that exerts the following force and torque on the head of the filament:

$$\mathbf{F}_{\rm ext} = -\xi^{\rm H}_{\rm T}\dot{\mathbf{r}} \tag{15}$$

$$\mathbf{T}_{\rm ext} = -\xi^{\rm H}_{\rm R}(\mathbf{t}\times\dot{\mathbf{t}}) \tag{16}$$

$\xi^{\rm H}_{\rm T}$ and $\xi^{\rm H}_{\rm R}$ are the effective translational and rotational drag coefficients, respectively. They are treated as independent parameters of the model. $\dot{\mathbf{r}}$ is the velocity of the head, given by

$$\dot{\mathbf{r}} = \frac{(\kappa\theta'\theta'' - f + T')\,\mathbf{t}}{\xi_\parallel} + \frac{(-\kappa\theta''' + \theta' T)\,\mathbf{n}}{\xi_\perp}, \tag{17}$$

Force and torque balance for the head read

$$\mathbf{F}(0,t) + \mathbf{F}_{\rm ext}(t) = \mathbf{0} \tag{18}$$

$$\mathbf{M}(0,t) + \mathbf{T}_{\rm ext}(t) = \mathbf{0} \tag{19}$$

where $\mathbf{F} = T\mathbf{t} - \kappa\theta''\mathbf{n}$ and $\mathbf{M} = \kappa\theta'\mathbf{t}\times\mathbf{n}$ are the internal force resultant and bending moment defined in section I. Substituting those expressions in (18)-(19) and using (15)-(17) yields

$$T - \frac{\xi^H_T}{\xi_\parallel}(\kappa\theta'\theta'' - f + T') = 0 \tag{20}$$

$$-\kappa\theta'' - \frac{\xi^H_T}{\xi_\perp}(-\kappa\theta''' + T\theta') = 0 \tag{21}$$

$$\kappa\theta' - \xi^H_R\dot{\theta} = 0 \tag{22}$$

To obtain the scaled (dimensionless) forms analyzed in the paper, we scale lengths by $\ell$, active force densities by $\kappa/\ell^3$, and tension by $\kappa/\ell^2$. This yields

$$\zeta\theta''' - \theta'' - \zeta T\theta' = 0, \tag{23}$$

$$\zeta\gamma T' - T - \zeta\gamma\,(\beta - \theta'\theta'') = 0, \tag{24}$$

$$\theta' - \zeta_R\dot{\theta} = 0. \tag{25}$$

where $\gamma = \xi_\perp/\xi_\parallel$, $\zeta = (\xi^{\rm H}_{\rm T}/\xi_\perp\ell)$, and $\zeta_R = (\xi^{\rm H}_{\rm R}/\ell^3\xi_\perp)$.

#### III. NUMERICAL SCHEME

##### A. Linear stability and the eigenvalue problem

The eigenvalue problems of sections 3.1 and 4.1 are solved on a uniform grid of $N = 500$ interior points plus two ghost points on each side. The spatial derivatives are approximated by second-order-accurate central finite differences. The four boundary equations are then used to eliminate the four ghost variables. The resulting discrete eigenvalue problem is solved numerically.

The finite difference formulas for the bulk equations are:

$$D^{(1)}_{\rm ij}\Theta_{\rm j} = \frac{\Theta_{\rm i+1}-\Theta_{\rm i-1}}{2(ds)}, \tag{26}$$

$$D^{(2)}_{\rm ij}\Theta_{\rm j} = \frac{\Theta_{\rm i+1}-2\Theta_{\rm i}+\Theta_{\rm i-1}}{2(ds)^2}, \tag{27}$$

$$D^{(3)}_{\rm ij}\Theta_{\rm j} = \frac{\Theta_{\rm i+2}-2\Theta_{\rm i+1}+2\Theta_{\rm i-1}-\Theta_{\rm i-2}}{2\,(ds)^3}, \tag{28}$$

$$D^{(4)}_{\rm ij}\Theta_{\rm j} = \frac{\Theta_{\rm i+2}-4\Theta_{\rm i+1}+6\Theta_{\rm i}-4\Theta_{\rm i-1}+\Theta_{\rm i-2}}{(ds)^4}, \tag{29}$$

where $ds = 1/(N-1)$ is the grid size. For example, the discretized bulk equation in the pinned-free and clamped-free cases is:

$$\Omega\Theta_{\rm i} = -D^{(4)}_{\rm ij}\Theta_{\rm j} + \beta(s_{\rm i}-1)D^{(2)}_{\rm ij}\Theta_{\rm j} + \beta D^{(1)}_{\rm ij}\Theta_{\rm j}. \tag{30}$$

Using the same formulas for the boundary conditions can make the ghost variable elimination singular. To avoid this problem we use modified formulas at orders 0, 1, and 2 that involve both ghost variables while remaining second-order-accurate and central:

$$D^{(0)}_{\rm ij}\Theta_{\rm j} = \frac{-\Theta_{\rm i+2}+4\Theta_{\rm i+1}+4\Theta_{\rm i-1}-\Theta_{\rm i-2}}{6}, \tag{31}$$

$$D^{(1)}_{\rm ij}\Theta_{\rm j} = \frac{-\Theta_{\rm i+2}+8\Theta_{\rm i+1}-8\Theta_{\rm i-1}+\Theta_{\rm i-2}}{12\,(ds)}, \tag{32}$$

$$D^{(2)}_{\rm ij}\Theta_{\rm j} = \frac{-\Theta_{\rm i+2}+16\Theta_{\rm i+1}-36\Theta_{\rm i}+16\Theta_{\rm i-1}-\Theta_{\rm i-2}}{12\,(ds)^2}. \tag{33}$$

#### B. Non-linear solutions by time stepper methods

Nonlinear solutions are obtained by discretizing the equations of motion and boundary conditions in space and time. The spatial derivatives are approximated by second-order-accurate finite difference approximations on a uniform grid with $N = 501$ points. This time there are not ghost points; instead we use asymmetric finite difference formulas near the boundaries. The equation for the tension $T(s,t)$, which has no explicit time derivatives, is solved independently at each time step, then fed into the equation for the angle $\theta(s,t)$. Time integration uses the second-order-accurate implicit-explicit scheme proposed by Tornberg and Shelley[1], wherein the fourth order spatial derivative is treated implicitly while the other (nonlinear) terms are treated explicitly:

$$\frac{3\theta^{n+1}_{\rm i}-4\theta^{n}_{\rm i}+\theta^{n-1}_{\rm i}}{2(dt)} = -D^{(4)}_{\rm ij}\theta^{n+1}_{\rm j} + 2G^{n}_{\rm i} - G^{n-1}_{\rm i} \tag{34}$$

$$G^{n}_{\rm i} = D^{(1)}_{\rm ij}\theta_{\rm j}D^{(1)}_{\rm ik}T_{\rm k} + T_{\rm i}D^{(2)}_{\rm ij}\theta_{\rm j} + \Delta_{\rm i} \tag{35}$$

$$\Delta_{\rm i} = \gamma\, D^{(1)}_{\rm ij}\theta_{\rm j}\left(D^{(1)}_{\rm ij}T_{\rm j} + D^{(1)}_{\rm ij}\theta_{\rm j}D^{(2)}_{\rm ik}\theta_{\rm k} - \beta\right) \tag{36}$$

where the superscript indicates the time step. $dt$ was varied between $2\times10^{-9}$ and $10^{-5}$ depending primarily on the value of $\beta$.

### SI-MOVIES

#### ESM Movie 1: Rotating state

Dynamics of a filament-cargo assembly as obtained by integrating the nonlinear equations of motion with $\gamma = 2$, $\zeta = 1$, $\zeta_{\rm R} = 10^{-3}$, and $\beta = 200$ corresponding to the top right corner of Figure 6b in the main text. The red circle indicates the head. Its size and shape are arbitrary. The black line is the trajectory of its center.

##### ESM Movie 2: Beating state

Same as movie 1 except $\zeta_{\mathrm{R}} = 10^{-1}$ corresponding to the top right corner of Figure 6b in the main text.

##### ESM Movie 3: Alternating state

Same as movie 1 except $\zeta_{\mathrm{R}} = 2 \times 10^{-3}$ corresponding to a point near the red/blue transition on the right edge of Figure 6b in the main text. The filament periodically alternates between rotating and beating. The exact trajectory is quite sensitive to the timing of the changes of states thus slight changes to the parameters' values may yield a fairly different looking trajectory.

##### ESM Movie 4: Effect of weak noise

The movie shows the dynamical trajectory of three swimmers starting at the origin where the active forces are always aligned along the local tangent. For the lowest force ($F = 5$), the swimmer moves slowly in almost straight trajectories with minimal bending. As the force density is increased ($F = 11$), the swimmer exhibits rotating modes of locomotion - here, the rotations (with the head tracing a circle or part thereof) persists for a while before switching direction due to the accumulated effects of noise. In the absence of noise this would correspond to the purely rotating mode with the head executing a closed orbit. For higher force densities, the swimmer exhibits flapping motion with persistent gait. All three swimmers have the same length and head size as seen from the number of beads. Simulations parameters are $\mathcal{K} = 20$, $\gamma = 1$ and $\kappa_{\mathrm{E}}\sigma^2/k_{\mathrm{B}}T = 20000$.

[1] Anna-Karin Tornberg and Michael J. Shelley. Simulating the dynamics and interactions of flexible fibers in stokes flows. *Journal of Computational Physics*, 196(1):8–40, May 2004. ISSN 0021-9991. doi:10.1016/j.jcp.2003.10.017.
